# Functional Electrical Stimulation and Brain-Machine Interfaces for Simultaneous Control of Wrist and Finger Flexion

**DOI:** 10.1101/2024.08.11.607263

**Authors:** Matthew J. Mender, Ayobami L. Ward, Luis H. Cubillos, Madison M. Kelberman, Joseph T. Costello, Hisham Temmar, Dylan M. Wallace, Edanjen T. Lin, Jordan L. W. Lam, Matthew S. Willsey, Nishant Ganesh Kumar, Theodore A. Kung, Parag G. Patil, Cynthia A. Chestek

## Abstract

Brain-machine interface (BMI) controlled functional electrical stimulation (FES) is a promising treatment to restore hand movements to people with cervical spinal cord injury. Recent intracortical BMIs have shown unprecedented successes in decoding user intentions, however the hand movements restored by FES have largely been limited to predetermined grasps. Restoring dexterous hand movements will require continuous control of many biomechanically linked degrees-of-freedom in the hand, such as wrist and finger flexion, that would form the basis of those movements. Here we investigate the ability to restore simultaneous wrist and finger flexion, which would enable grasping with a controlled hand posture and assist in manipulating objects once grasped. We demonstrate that intramuscular FES can enable monkeys with temporarily paralyzed hands to move their fingers and wrist across a functional range of motion, spanning an average 88.6 degrees at the metacarpophalangeal joint flexion and 71.3 degrees of wrist flexion, and intramuscular FES can control both joints simultaneously in a real-time task. Additionally, we demonstrate a monkey using an intracortical BMI to control the wrist and finger flexion in a virtual hand, both before and after the hand is temporarily paralyzed, even achieving success rates and acquisition times equivalent to able-bodied control with BMI control after temporary paralysis in two sessions. Together, this outlines a method using an artificial brain-to-body interface that could restore continuous wrist and finger movements after spinal cord injury.

**One Sentence Summary:** We show in non-human primates that intramuscular functional electrical stimulation can flex the wrist and fingers through a functional range of movement, can be controlled precisely, and that a brain-machine interface can determine command signals for intended finger and wrist flexion.

## INTRODUCTION

Spinal cord injuries (SCI) impact a person’s ability to perform activities of daily living and negatively impact quality of life, especially when hand function is affected. Almost 60% of SCI cases in the US are partial or complete injuries at the cervical level which typically decreases hand and arm function (*1*). Restoration of hand function is a top priority for people with spinal cord injuries to improve quality of life (*2, 3*). Functional electrical stimulation (FES) can restore hand movements after severe spinal cord injuries by electrically activating a person’s own muscles. Stimulating electrodes are typically placed either at the nerves proximal to the muscles (epineural or intrafascicular) or at the muscles themselves. FES has been used to restore hand movements since the 1980s (*4*) and prescribed movements such as lateral and palmar grasps have been restored in more than 220 users (*5*). For restoring natural hand control beyond prescribed grasps, it is necessary to continuously control movements in many degrees-of-freedom (DOF) of the hand simultaneously. For example, functional tasks may require grasping to different degrees, i.e. for grasping different sized objects, or in different postures. In particular, activities of daily living require a broad range of wrist postures in addition to grasping (*6*). Controlling wrist flexion would allow users to stabilize the hand, reach new objects, augment the grasp itself (*7, 8*), and move the grasped object. However, clinical trials with FES thus far have only focused on restoring specific grasps (*9, 10*) and have yet to control wrist posture at the same time.

Restoring graded control to the wrist and fingers can be broken down into a control problem with two parts: first, estimating the intended movements of multiple-DOF in the hand to use as stimulation commands, and second, stimulating to restore multiple-DOF movements. With improving technologies and algorithms, brain-machine interfaces (BMIs) have recently reached unprecedented levels of success in estimating user intentions for various applications such as decoding speech (*11, 12*), controlling computer cursors (*13, 14*), and moving prosthetic arms (*15*). Whereas control schemes using residual movements (*5, 16, 17*) or muscle activations (*18–21*) can be unintuitive or not function in more severe injuries, using BMIs provides a control signal using an intact part of the natural motor control pathway (*9*). Intracortical BMIs in particular are promising for estimating graded intended movements of multiple-DOF in the hand. BMIs have been used to decode multiple-DOF of the upper limb at the same time during prehensile tasks (*15, 22*), and one or a few continuous DOF movements in the hand such as two finger groups (*23*), three finger groups with 4-DOF (*24*), and hand or wrist flexion (*25*). With respect to decoding finger and wrist movements, changes in wrist posture introduce errors to finger movement predictions (*26*). While predicting finger movements in new wrist postures is still achievable (*26, 27*), simultaneously decoding continuous wrist and finger movements has yet to be demonstrated.

Still, BMI performance controlling animated hands has arguably outpaced the dexterity of movements that can actually be restored using FES. Fully restoring movements to both the wrist and fingers at the same time is difficult. These joints are biomechanically linked, with the muscles that control the fingers crossing the wrist joint, both changing length with wrist posture and evoking moments about the wrist when moving fingers. To avoid these difficulties, previous FES studies have instead grouped the wrist and fingers (*28*) or controlled the wrist and fingers at different times (*25*). Intramuscular FES is a promising approach to restore the multiple DOF required for dexterous hand movements. Whereas intrafascicular or nerve cuff stimulation can create functional grasps by evoking synergistic movements (*29, 30*), intramuscular FES allows targeting specific muscles or even portions of a muscle innervated by a single peripheral branch (neuromuscular compartments) (*31*). Stimulation at this most distal point more easily evokes selective movements at the expense of requiring more electrodes to cover all the target muscles. Additionally, stimulation can be modulated to evoke different sized movements (*32–34*), leading to graded control in many DOF. Intramuscular FES has been used to restore continuous finger opening and closing (*25, 35*), wrist torques (*36*), as well as discrete grasps (*10*); however, it is unknown how well intramuscular FES could restore continuous finger and wrist movements at the same time.

Here, we first evaluate how well we can continuously control movements of the wrist and hand using intramuscular FES. We show that we can evoke flexion movements in either the wrist or the fingers by stimulating muscles for each DOF independently, moving the fingers through a range of flexion larger than previous FES studies (*10, 37*) and reported functional ranges during activities of daily living (*38, 39*), while moving the wrist through a range of motion equivalent to the functional range of motion in activities of daily living (*6, 40*). Then we show that by combining stimulation for both DOF we can evoke similarly sized movements while also controlling wrist posture, and, for the first time to our knowledge, reach target finger and wrist flexion simultaneously. We also quantify the effects of fatigue and the interaction between the two DOF. We then demonstrate a BMI for predicting intended wrist and finger movements in real-time. We show that a linear 2-DOF BMI works well, and while the BMI control efficacy decreases after blocking sensory afferents, we find that performance can be recovered by retraining the model, even matching able-bodied task performance in a subset of sessions.

## RESULTS

### Experiment overview

Two rhesus macaque monkeys (Monkey R and Monkey N) performed a virtual target acquisition task in which the goal was to move the virtual hand to target levels of finger and wrist flexion (Figure 1A). Both monkeys performed this with target-controlled FES during which the monkey’s hand was temporarily paralyzed via pharmacological nerve block and intramuscular stimulation moved the monkey’s hand to acquire targets with the virtual hand mimicking their hand (Figure 1A, orange). Stimulation was delivered through chronic bipolar intramuscular electrodes implanted in muscles of the forearm that control the hand and wrist (Figure 1B). A list of electrode targets is included in Supplemental Table 1. Monkey N also performed the task with hand control, in which they moved their able-bodied hand in a manipulandum to control the virtual hand (Figure 1A pink), and brain-machine interface (BMI) control, in which their decoded intended movements directly controlled the virtual hand (Figure 1A blue). Monkey N was previously implanted with Utah electrode arrays in the hand area of primary motor cortex (Figure 1C) which provided neural signals for BMI control. Evoked movements from stimulation on intramuscular electrodes were also tested outside of the virtual target acquisition task with movements characterized by video and either manual annotation or Deeplabcut (*41, 42*) (Figure 1D).

**Fig. 1.**
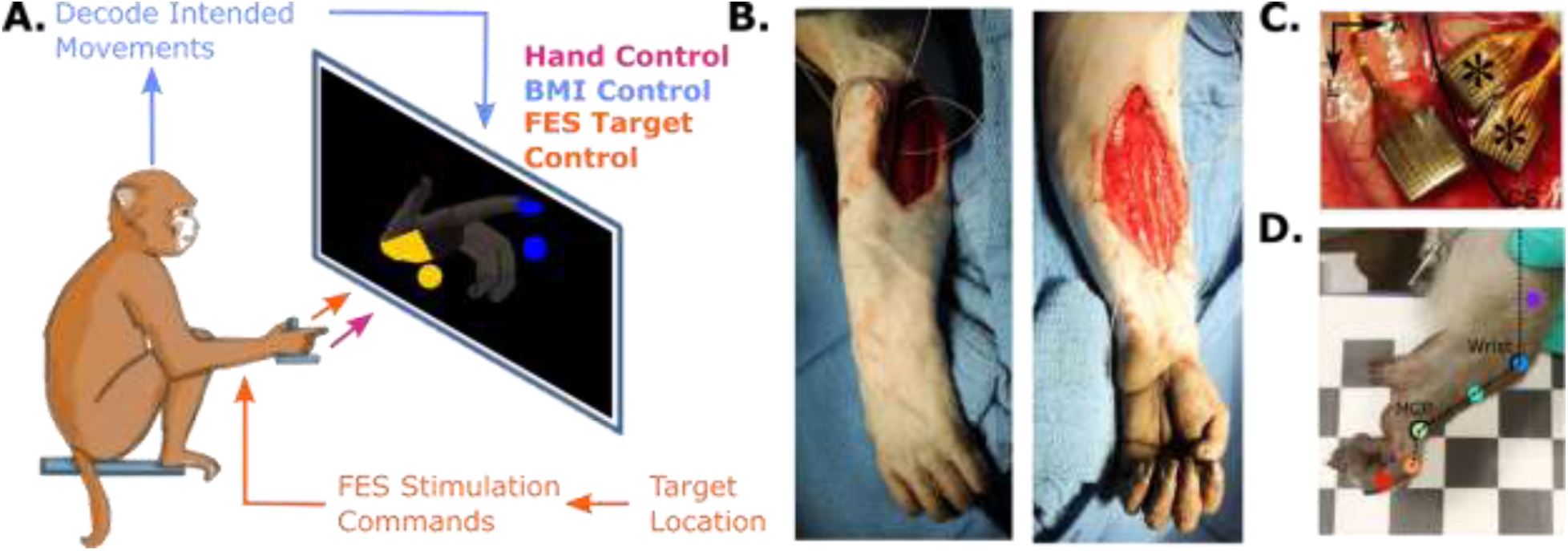
Experimental setup. (**A**) Illustration of the three different types of closed-loop experiments performed: hand control, BMI control, and FES target control. (**B**) Intraoperative images from intramuscular electrode implantation in Monkey R. Electrodes were implanted in flexor and extensor muscles via either the dorsal (left) or volar (right) surgical incision and then tunneled across the shoulder to a percutaneous exit site on the back. (**C**) Location of Utah array implants in Monkey N. The two arrays marked with (*) are anterior to the central sulcus, in the primary motor cortex, and were used in this study (CS – central sulcus, A – anterior, L – lateral). (**D**) Example angle measurements using markers identified with Deeplabcut (*41, 42*). Example wrist flexion and finger flexion (MCP - metacarpophalangeal) angles are marked.

### Coordinating electrode stimulation with patterns

A benefit of intramuscular stimulation is that stimulation on individual electrodes is expected to activate portions of individual muscles and therefore evoke more selective movements which can then be combined into functional movements. We first asked how well we can control the wrist and finger DOFs individually using coordinated stimulation of multiple intramuscular FES electrodes. Following conventional FES control methods (*43*), a stimulation pattern was defined for each of the two DOF (wrist and finger flexion). These patterns define stimulation parameters that extend the desired DOF at low commands and flex the desired DOF at high commands. Patterns were established using initial single electrode stimulation results, as described in the Methods, then minimally adjusted on each day. Supplemental Figures 1A and 1B show example finger and wrist patterns respectively for Monkey R. We observed that each pattern successfully moved the intended DOF across a large range of motion (ROM) (Figures 2A and 2B).

**Fig. 2.**
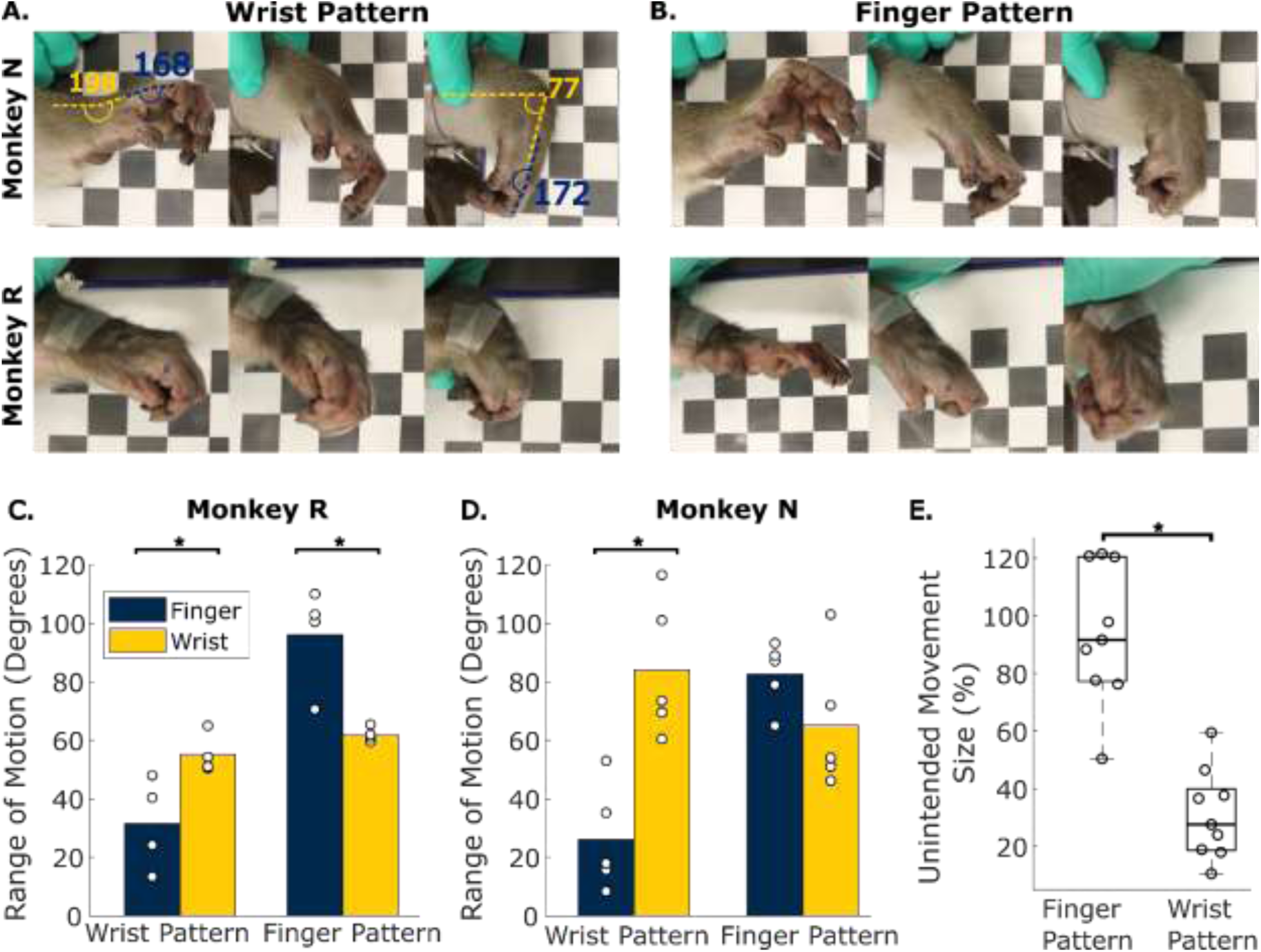
Range of motion using stimulation parameters independently. (**A**) Example evoked movements from testing the wrist stimulation pattern on one example day with each monkey (top – Monkey N, bottom – Monkey R). (**B**) Example evoked movements from testing the finger stimulation pattern on one example day with each monkey (top – Monkey N, bottom – Monkey R). (**C-D**) The range of evoked movements in the index MCP joint (blue) and wrist joint (yellow) using the wrist pattern and finger pattern for Monkey R (C) and Monkey N (D) respectively. Open circles indicate measurements from different experiments where patterns were tested. (**E**) Size of “unintended” movements, measured as the size of the movement on the opposite joint, i.e. wrist joint for finger patterns and finger joint for wrist patterns, presented as a percentage of the ROM given by the intended pattern for that joint. Open circles indicate measurements from different experiments. Results are grouped for both monkeys. Boxes indicate the 25^th^ to 75^th^ percentiles and the central mark indicates the median.

To quantify these movements, we measured the joint flexion angles for the wrist joint and the index metacarpophalangeal (MCP) joint during each evoked movement. Supplemental Figures 2A and 2B illustrate example measurements for one session testing commands throughout the wrist and finger pattern respectively. When tested in multiple sessions, the finger patterns moved the fingers through a large ROM for both monkeys (Figures 2C and 2D), averaging 88.6 degrees (std=15.0) pooled across both monkeys. Similarly, the wrist patterns moved the wrist through an average range of 71.3 degrees (std=23.0) pooled across both monkeys, which is not significantly different from the 80 degree functional wrist flexion range reported in studies of activities of daily living (*6*) (p=0.29, one-sample t-test). Additionally, the evoked finger movements are larger than reported functional MCP flexion ranges of 53.3 degrees and 52 degrees (*38, 39*) and the approximately 40 to 50 degree range in MCP previously reported to be restored with intramuscular FES (*10*).

While stimulation achieved large ranges of motion, both patterns also evoked unintended movements in the other joints (Figure 2E). Most notably, the finger patterns for both monkeys evoked large wrist movements, with a median range 91.7% the size of the wrist ROM evoked by the wrist patterns, likely due to these muscles crossing the wrist joint. These unintended movements indicate that it would be difficult to individuate the fingers from the wrist using only the finger pattern. Wrist patterns, however, evoked less unintended movements, moving the fingers through a median range only 27.9% the size of finger ROM evoked by finger patterns. Additionally, wrist patterns moved the wrist joint in a range 150.4% larger than they moved the fingers (p<0.01 paired t-test). The relative selectivity with which wrist patterns can evoke wrist movements suggests that we may be able to individuate the wrist from the fingers by using the wrist pattern to stabilize the wrist during finger movements.

### Simultaneous stimulation on both patterns can move both DOF separately

We next tested whether, by using these two patterns simultaneously, we could successfully move the wrist and fingers separately, i.e. opening or closing the fingers while using wrist stimulation to maintain wrist flexion or extension. Figure 3A and 3B show example evoked movements during these tests for Monkey R and Monkey N respectively. We measured the finger and wrist angles during these four movements within an experimental session to illustrate the ROM for each session (Figures 3C and 3D). Measured angles were calibrated to the range of motion measured during individual pattern stimulation, as described in the methods. For example, a wrist extension of 1 is the angle that the wrist pattern extended the wrist to on that day and a finger extension of 0 is the angle that the finger pattern flexed the fingers to on that day. When testing these four postures, evoked movements explored a large portion of the 2-DOF ROM, averaging 101.8% (std = 51.9%) the size of the calibration range.

**Fig. 3.**
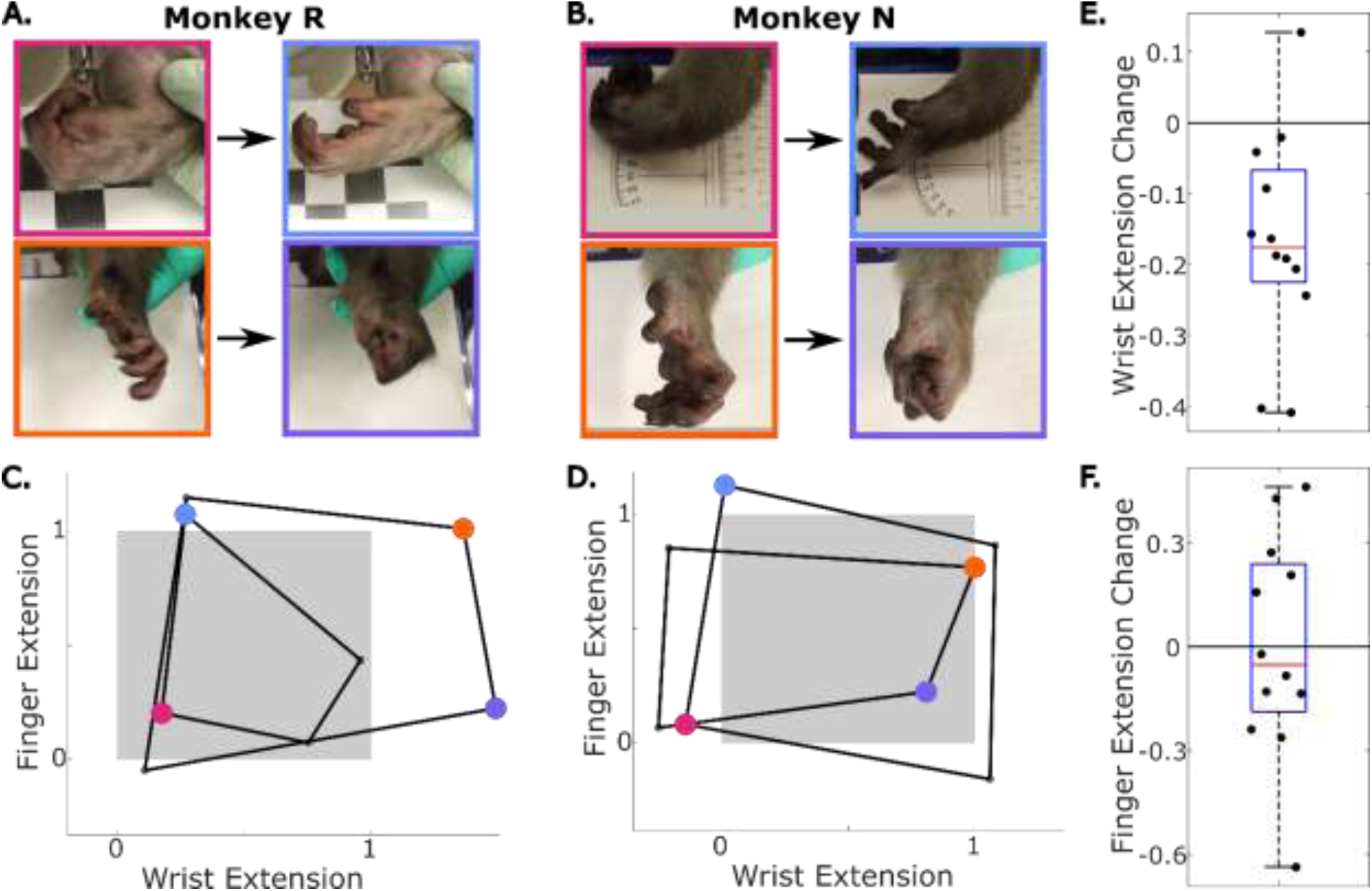
Range of motion using two-patterns simultaneously. (**A**) Example stimulated movements opening or closing the fingers with simultaneous stimulation holding the wrist in flexion (top) or extension (bottom). Results for Monkey R. (**B**) Same as (A) but for Monkey N. Measured finger and wrist angles when opening and closing the fingers with the wrist held fully extended or fully flexed. Angles for postures on the same day are connected to make one quadrilateral. All four postures were tested on two days for each monkey. Colored points correspond to the frames in (A) with matching colored borders. Extensions are calibrated to the 1-DOF ROM available (Methods). The grey box is a unit square corresponding to the calibration. Same as (C) but for Monkey N. (**E**) Change in wrist extension (fraction of calibrated ROM) when the fingers are stimulated to flexion with stimulation holding the wrist either flexed or extended. This was done on four days for Monkey N and three days for Monkey R. (**F**) Change in finger extension (fraction of calibrated ROM) when the wrist is being held extended instead of flexed.

There are two effects due to biomechanical coupling that we expect to impact the ROM available in 2-DOF stimulation. First is active coupling, where extrinsic finger muscles exert torque on the wrist when activated. The second is passive coupling, where changes in wrist posture change the tendon length of extrinsic finger muscles making finger flexion or extension easier. We found a large active effect as flexing the fingers also significantly flexed the wrist with a median change in wrist extension of -17.5% of the movement range (p=0.0015) (Figure 3E). Alternatively, the fingers tended to be more flexed at wrist extension, but the effect was not significant (Figure 3F). These results suggest that active effects should be heavily considered when designing control schemes, while passive effects have much less impact.

Having defined the limits of the movement range explored by simultaneous stimulation, we next tested how well stimulation evoked movements in both DOF throughout the 2-DOF stimulation space. Figure 4A illustrates example movements for each monkey as stimulation was stepped between random points along the two patterns. As might be expected, we found that pattern command levels could predict the evoked postures of their intended DOF with some accuracy (Figure 4B). In one example session with each monkey, finger pattern stimulations predicted evoked finger posture with an R^2^ of 0.39 and 0.77 for Monkey R and N respectively (Figure 4B top), while wrist pattern stimulation predicted wrist posture with an R^2^ of 0.20 and 0.45 (Figure 4B bottom). There was, however, a lot of variance in the evoked posture that was not explained by stimulation on only one pattern. Finger pattern commands also explained a significant amount of evoked wrist posture, with a R^2^ of 0.35 and 0.12 for Monkey R and Monkey N respectively (Figure 4C). As a result, wrist postures were predicted better using stimulation information from both patterns compared to only using wrist stimulation, a 0.43 and 0.10 increase in R^2^ for Monkey R and Monkey N respectively. This effect was smaller for the finger postures which are expected to have no active effect from wrist muscles. Using both stimulation parameters to predict finger angle resulted in an R^2^ increase of 0.09 over using only finger stimulation for Monkey R, and no significant change for Monkey N. This indicates that stimulation control schemes will need to account for the active torque on the wrist due to finger muscle stimulation throughout the 2-DOF ROM as well and the effect of wrist posture on finger muscles is less of a consideration.

**Fig. 4.**
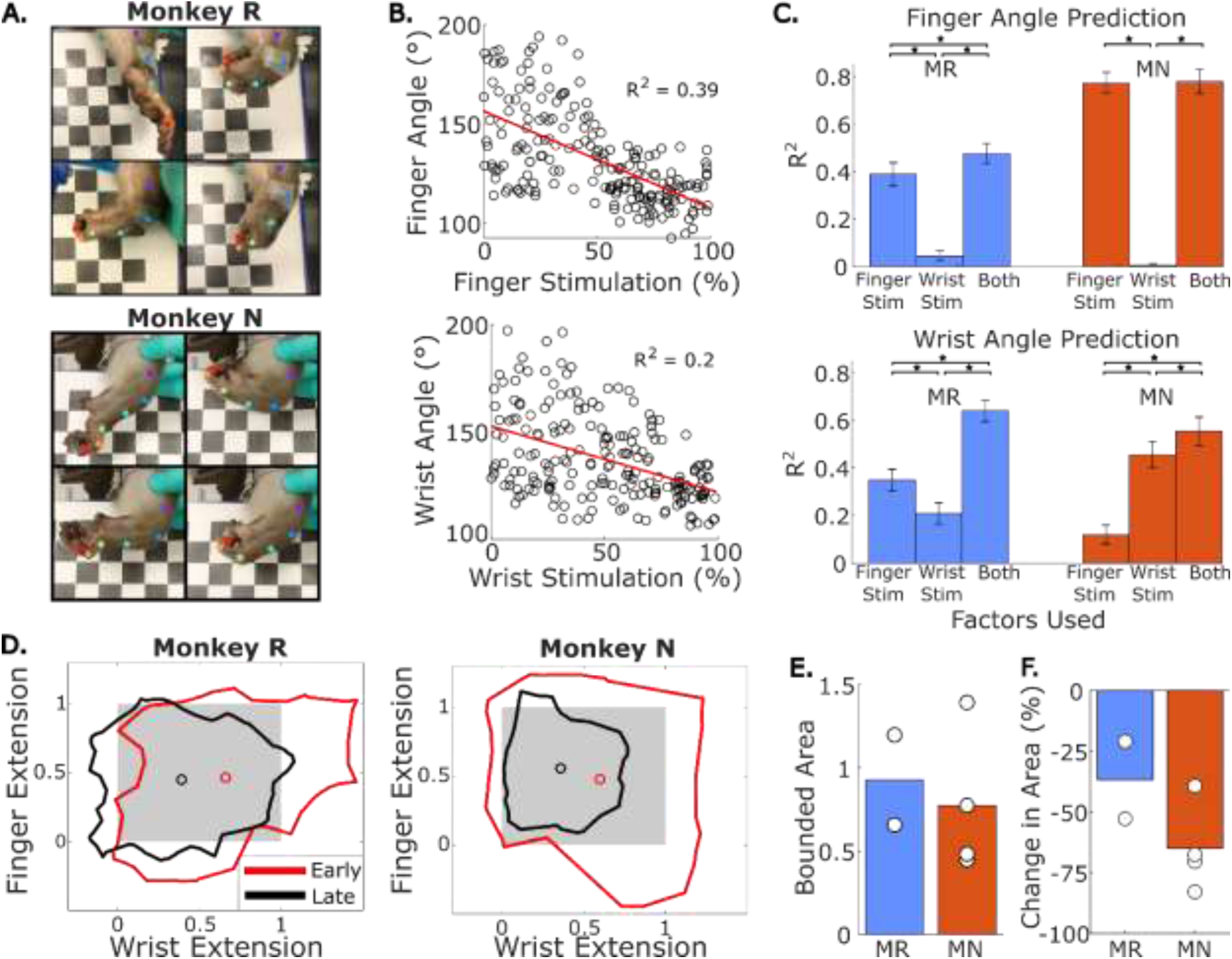
Movement range using graded stimulation on both patterns. (A) Example frames from video of 2-DOF stimulation with markers estimated using DLC for Monkey R (top) and Monkey N (Bottom). (B) Evoked posture versus commanded stimulation intensity for the pattern intended to move that joint (top – finger, bottom – wrist). Angles are taken as the average joint angle during the last 0.5 seconds of the 3 second stimulation. Red lines indicate linear fits between stimulation intensity and joint angle. (C) Variance explained in finger angles (top) and wrist angles (bottom) by the linear fits from the factors given by the x-axis, either finger pattern stimulation command, wrist pattern stimulation command, or both. MR = Monkey R. MN = Monkey N. (D) Boundaries drawn containing measurements of each finger extension and wrist extension from video frames when stimulation was being administered in the first 3 minutes (red) or last 3 minutes (black).

In testing 2-DOF ROM during the randomly-sampled stimulation experiments, we observed noticeable changes in the ROM over the course of each experiment. Stimulation sessions lasted for several minutes (average 8.35 minutes [std=1.33] or 167 stimulation trials [std=26.6]), and we observed a decrease in the evoked ROM over the course of the experiment. Figure 4D illustrates the evoked ROM for each monkey. The enclosed area indicates a boundary on the measured movements in each joint with the red boundary indicating trials in the first three minutes (approximately 60 trials) and the black boundary indicating later trials in the last three minutes. We observed initial movements through a large portion of the finger and wrist movement space for both monkeys that diminished over time, presumably due to fatigue. In the first three minutes, Monkey R’s movement range area averaged 0.93 and Monkey N’s movement range area averaged 0.77 (Figure 4E). In the last three minutes, Monkey R’s movement area was 36.9% smaller and Monkey N’s movement area was 65.1% smaller than movements in the first three minutes (Figure 4F). This fatigue effect was largest for wrist extension for both monkeys (Supplemental Figure 3), which can be seen by the centroid of the bounded area shifting away from wrist extension in late trials.

Angles are pooled from two days with Monkey R and four days with Monkey N. Angles are calibrated to the ROM available from stimulation in the respective pattern for that day. The frames at the highest and lowest 0.1% of measurements in each DOF are excluded. Grey box is a unit square corresponding to the calibration. (E) Size of bounded area from early trials. Area is calculated using calibrated movements so that values indicate a proportion of the calibrated ROM. Each dot indicates the measurement for one experimental day, there were two days for Monkey R and four days for Monkey N. (F) Change in bounded area between the first 3 minutes and last 3 minutes using two days for Monkey R and four days for Monkey N.

### Continuous control of 2-DOF FES to reach target flexion

With a large range of 2-DOF wrist and finger movements available using FES, it is desirable to continuously control this stimulation to reach specific postures throughout the available ROM. To test this, we used a virtual finger and wrist target acquisition task in which targets were presented on a screen and then used FES to move the monkey’s nerve-blocked hand to acquire those targets (see Figure 5a for an example in Monkey R). To enable targets to be reached at even moderate levels of fatigue, targets were restricted to a reduced range consisting of the middle 50% of the ROM for each joint. In each experiment we obtained greater than 84% success (Figure 5D). However, we observed that both monkeys had difficulty completing trials where the wrist was extended and the fingers were flexed (Figures 5B and 5C). In the most extreme wrist extension/finger extension splits, Monkeys R and N only achieved 50% and 33.3% rates respectively. This is likely due to FDP activation during finger flexion, which is known to also produce some wrist flexion. While success rates were high, average acquisition times were much slower than a representative able-bodied control acquisition time from Monkey N (Figure 5E). Notably, FES control was only investigated with slow updates (i.e. stepping one command value out of 255 every 32ms) with the intention of reducing oscillation near targets. However, this continuous control shows enough fine control with FES to reach precise postures within a reduced range.

**Fig. 5.**
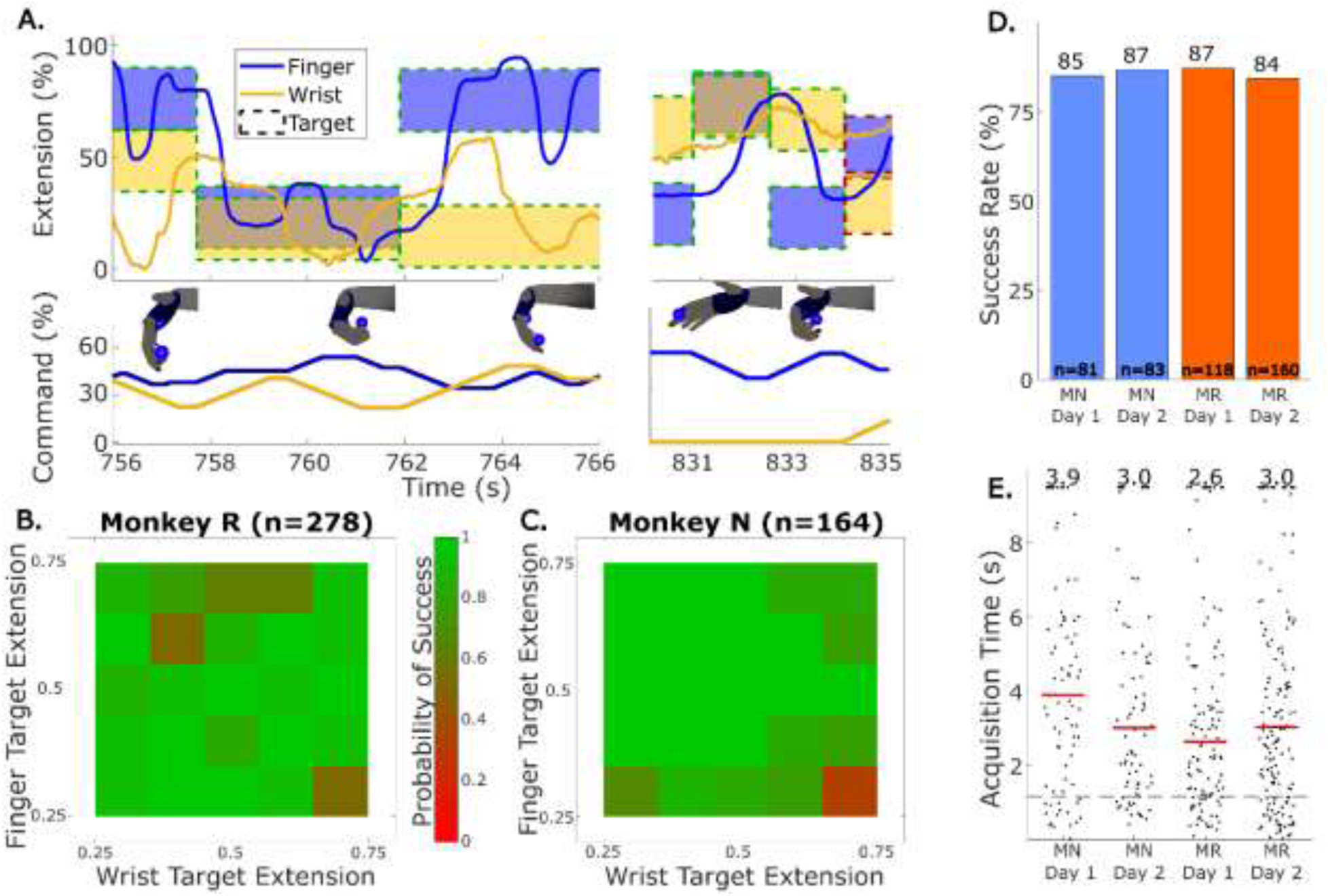
Closed-Loop control of two-DOF FES. (**A**) Example finger and wrist extension (top) and FES commands (bottom) from closed-loop FES control. Blue and yellow patches indicate targets; a green border indicates success while a red border indicates failure. Representative images of the virtual hand are included to illustrate the target movement at that time. (**B**) Probability of closed-loop FES trial success by target location, pooled across two experiments for Monkey R. (**C**) Same as (B) but for Monkey N. (**D**) Success rate for closed-loop FES in each session, two for each monkey. MN=Monkey N, MR=Monkey R. (**E**) Acquisition times for successful trials using FES. Red bars and numbers above each column indicate the median acquisition time for that session. Dashed line indicates an example acquisition time from Monkey N using hand control.

### BMI control of virtual finger and wrist movements

We next asked how well we can generate a command signal from brain activity to ultimately control this 2-DOF FES aimed at restoring coordinated finger and wrist flexion movements. We trained a recalibrated feedback intention-trained (ReFIT) Kalman filter model (RKF), following the procedure we have used previously (*23, 26*), enabling Monkey N to control finger and wrist flexion of a virtual hand with only their activity from implanted Utah arrays (Figure 1A, BMI control). While these experiments were performed with Monkey N they were separated in time from FES due to a decline in brain signal quality. Monkey N was able to achieve high levels of success simultaneously controlling the wrist and finger flexion with a BMI. Figure 6A shows example decoded finger and wrist flexion from a real-time BMI session with Monkey N using the RKF model to quickly acquire targets. The monkey acquired targets with 97% success with a median acquisition time of 1.0s during this session. We tested 2-DOF BMI control of the wrist and finger DOF on three experimental days. Performance was measured with acquisition time, orbit time, and success rate (Figures 6B to 6D). Performance with the RKF model was comparable to that of using the hand to control the manipulandum with a similar success rate across sessions (96.5% down to 94.5%, p=0.041 chi-squared test), only one session with a higher median orbit time (311.5 ms increase, p<6e-8 two-sample Kolmogorov-Smirnov test), and two sessions showing increases in median acquisition time (391 ms and 489 ms increases, p<5e-11, two-sample Kolmogorov-Smirnov test). Due to the high performance we concluded that there is enough information in our array recording in the hand area to decode command signals for 2-DOF in real time with a linear Kalman filter.

**Fig. 6.**
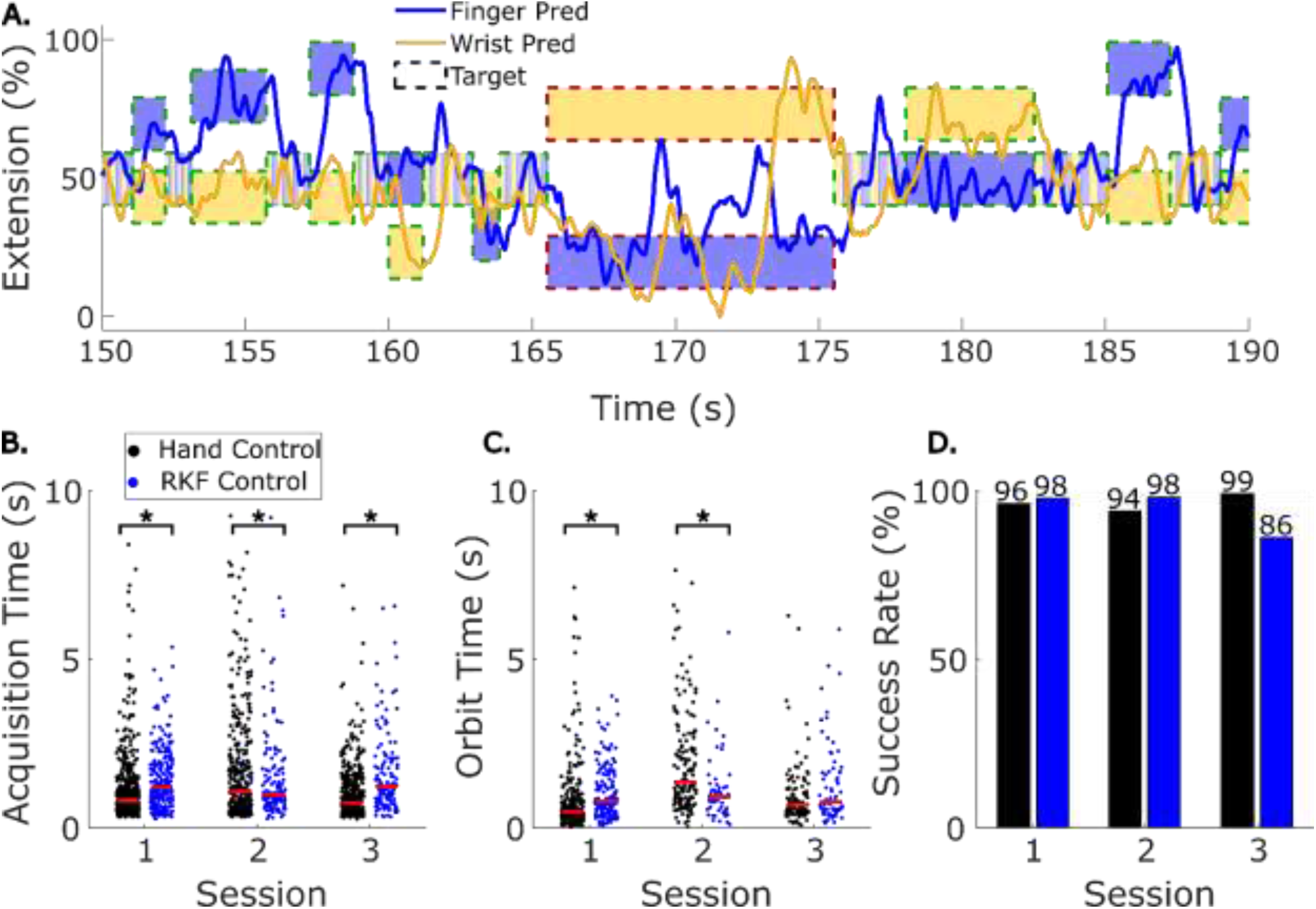
Two-DOF BMI control of virtual wrist and fingers. (**A**) Example decoded finger and wrist extension from a closed-loop BMI experiment. Green target boundary indicates trial success, and red target boundary indicates failure. (**B**) Distributions of acquisition time for all 3 sessions with RKF control in blue and hand control in black. Each dot indicates an individual successful trial. Red bars indicate the median. (**C**) Distribution of orbit times for all three sessions, following the format in (B). (**D**) Success rate using hand control (black) or BMI control (blue) during each session.

For translational purposes, it will be important to decode intended movements in a patient with spinal cord injury whereas our monkey model is able-bodied. To this end, we tested whether the trained Kalman filter could still decode two-DOF of intended movements in a real-time BMI task after the same nerve block used in FES experiments, which eliminates movement and proprioception similar to a spinal cord injury. First, an RKF model was trained and tested prior to the nerve block. Then the nerve block was performed and the same RKF model was tested again (Figure 7A). Figure 7B and 7D show average acquisition time and success rate respectively during each set of trials in one example session. In this example session, BMI performance decreased while using the same RKF model after nerve block, median acquisition time increased from 971ms to 1611ms (p<1e-5, two-sample Kolmogorov-Smirnov test), and success rate decreased from 98% to 93% (p=0.03, chi-squared test).

**Fig. 7.**
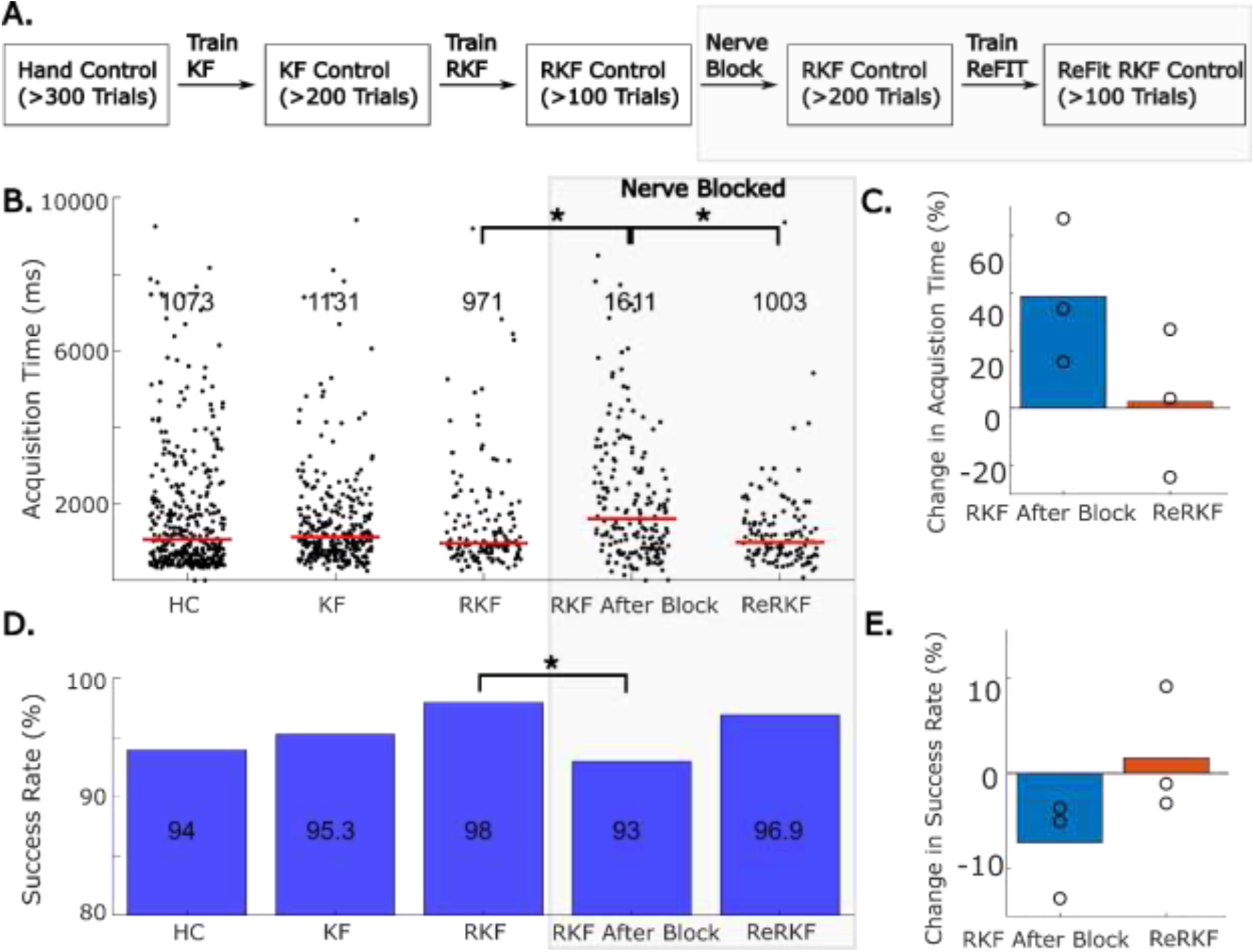
Recovering BMI performance after nerve block. (**A**) Illustration of the model training process with respect to the nerve block. (**B**) Acquisition times during an example session. ReRKF is the decoder that was trained in a second ReFIT step after nerve block. Red lines and numbers near the top of each column indicate the median acquisition time for that set of trials. (**C**) Change in median acquisition time during three experiments where an RKF was tested before and after nerve block and then compared to the ReRKF. The change is measured with respect to RKF performance before nerve block. (**D**) Success rates during an example session. (**E**) Percent change in success rate during the three experiments where RKF was tested before and after nerve block and compared to ReRKF. Change is measured with respect to RKF performance before nerve block.

Interestingly, some performance was regained by applying a second ReFIT calibration stage using the successful trials from BMI control after nerve block. In the example session, using the retrained RKF (ReRKF) improved median acquisition time to 1003 ms (from 1611 ms; p<4e-5, two-sample Kolmogorov-Smirnov test) and success rate to 96.9% (from 93%; p=0.13, chi-squared test). We repeated the BMI task in three nerve block experiments. First, when testing the same RKF before and after nerve block, BMI performance consistently decreased after the nerve block. Across these three days, acquisition time increased by an average of 38.8% (p<1e-9, two-way ANOVA) (Figure 7C) and success rate decreased by 7.2% (p<1e-4, chi-squared test) (Figure 7E). Then, retraining the RKF using ReFIT on BMI trials after the nerve block successfully regained performance in all three experiments. ReRKF performance showed equivalent acquisition times (p=0.32, two-way ANOVA) (Figure 7C) and success rates (p=0.68, chi-squared test) (Figure 7E) compared to before block RKF performance. Performance metrics for each experimental session are given in Supplemental Table 2. Notably, the success rates and acquisition times in the first two sessions were not significantly different between hand control trials and the ReRKF trials with a median acquisition time increasing 103ms (p=0.047, two-sample Kolmogorov-Smirnov test) in one session and decreasing 70ms (p=0.037, two-sample Kolmogorov-Smirnov test) in the other. Ultimately this shows that a linear Kalman filter can decode intended finger and wrist movements in 2-DOF after nerve block despite the loss of sensory information.

## DISCUSSION

In recent years, BMI studies have demonstrated unprecedented success in uses such as controlling computer tablets and cursors (*13, 14, 44*), controlling prosthetic arms (*45*), and decoding intended text and speech (*11, 12*). Controlling FES on independent DOF is analogous to BMI control in which independent joint angles are decoded from neural activity (*25, 35*). Previous studies have decoded finger movements (*23, 46*), or decoded wrist movements (*25*). Here we show that Utah microelectrode arrays implanted in finger area of primary motor cortex of one animal can decode simultaneous finger flexion and wrist flexion, achieving BMI performance similar to performance controlling the virtual hand with their able-bodied hand. To our knowledge, this is the first work showing continuous decoding of simultaneous finger and wrist movements, achieving performance near the monkey’s able-bodied control of the manipulandum and virtual hand. Additionally, we show that this decoding approach works after nerve block with a retraining step, indicating that online recalibration methods, similar to methods that have become standard in many BMI studies (*47–49*), work well in NHP after sensory afferents are lost.

While BMIs have demonstrated the ability to decode dexterous movements and continuous movements in multiple-DOF, previous FES studies have primarily only restored discrete functional grips (*10, 29, 37, 50*), one-DOF continuous finger movements (*35, 51*), or two-DOF continuous movements in the arm (*25, 28*). Intramuscular FES is promising for restoring dexterous hand movements due to its ability to selectively activate portions of muscles and evoke selective movements. With more movements available, stimulating with multiple electrodes can cover a larger basis of hand movements. Here we show an implementation of intramuscular FES to restore wrist and finger flexion independently which then formed the basis for successfully controlling continuous finger and wrist flexion simultaneously to achieve arbitrary postures. In addition, we characterized how well we could combine stimulation on two biomechanically linked DOF, the wrist and the fingers, simultaneously. We found that stimulation successfully evokes large movements, greater than the functional ROM for MCP joints during activities of daily living (*38, 39*) and similar to the wrists functional ROM (*6*). The evoked 2-DOF ROM is significantly impacted by interactions between evoked movements from stimulation on both patterns, primarily the large wrist movements caused by stimulating extrinsic finger muscles. Interestingly, the effect of wrist posture on evoked finger movements was much smaller.

Future work can aim to account for interactions between the DOF being controlled, both with respect to decoding accuracy and FES control. Ultimately, brain-controlled FES involves decoding an intended movement and a translation from intended movement to the stimulation parameters required to perform the movement. The successful BMI experiments with a linear model here indicate that the BMI algorithm could identify separate control signals for both the wrist and fingers, however, the FES evoked movements on the two-DOF are not independent, with wrist movements dependent on both wrist and finger stimulation. More work is needed to understand how well BMI systems could use increasingly successful neural network methods (*48, 52*), or feature engineering (*26, 53*), to better predict the required stimulation, or if the systems would be more effective when relying on the user to re-aim their BMI control (*26, 54, 55*).

With regards to stimulation protocols, we found that there was high variance in how well evoked movements matched the intended movement, as given by the stimulation command. The patterns relating stimulation to movement were designed by hand using single electrode characterizations and trial-and-error methods with little data to estimate movements from stimulation on combinations of electrodes. Alternative methods can design stimulation protocols more efficiently. For example, matching desired movements to EMG activity in multiple muscles and choosing stimulation to achieve those muscle activations uses able-bodied movements, which are easier to obtain, to design stimulation protocols (*56*). Alternatively, using algorithms such as Bayesian optimization has shown promise for efficiently tuning stimulation parameters and achieve desired movements (*57, 58*). These algorithms optimize an objective function related to the movement using stimulation on each electrode as an input, which deviates from current control methods where one electrode contributes to movements in one DOF.

Throughout these experiments muscle fatigue has been a limiting factor. In intramuscular FES, fatigue is caused by a few factors. Large motor units that fatigue quickly are recruited before or with smaller fatigue resistant fibers, a phenomenon known as reverse recruitment. Fibers are also recruited at relatively high rates. In normal recruitment, fiber activations are offset and a fused contraction is formed by many overlapping twitches offset in time. In FES, all of the fibers are activated at once, so a fused contraction must be created by stimulating at a higher rate. Here, significant fatigue was observed in some muscles after as few as three minutes of stimulation. More work is needed to integrate stimulation paradigms that can alleviate fatigue, such as using more electrodes to activate more of the muscle (*59*), interleaving stimulation on multiple electrodes (*60*), or targeting muscle motor points more specifically (*61*), into systems designed to restore selective and functional movements.

## MATERIALS AND METHODS

All procedures were approved by the University of Michigan Animal Care and Use Committee.

### Experimental design

Two male Rhesus Macaques (Monkey N and Monkey R) performed a virtual target acquisition task, moving their hands while finger and wrist posture were tracked. Monkey N did this task with hand control, BMI control, and target-guided FES control, whereas Monkey R only did this task with target-guided FES control. In hand control, the monkey’s hand moved a manipulandum to control the virtual hand. In FES target control, the monkey’s hand again moved the manipulandum to control the virtual hand, however the hand was temporarily paralyzed and moved due to stimulation. In BMI control, the monkey’s movement intentions decoded from neural activity moved a virtual hand. We used a computer running xPC Target version 2012b (Mathworks) to coordinate the experiment in real time, storing task parameters, neural data, and hand kinematics while coordinating target presentation, executing the decoder model to predict finger and wrist positions, and sending stimulation commands to the networked neuroprosthesis (NNP) (*35, 62, 63*). The NNP system then stimulated through the implanted electrodes. Both monkeys also underwent open-loop stimulation, in which electrodes were used for stimulation and hand movements were recorded with video but not displayed on the screen in front of the monkey. Open-loop stimulation was controlled either by the experimenter to test the range of the movements evoked with stimulation patterns (as in Figures 2 and 3), or by xPC to sample stimulations from the patterns (as in Figure 4).

### Implants

Both monkeys were implanted with chronic bipolar intramuscular electrodes (Synapse Biomedical, Inc, Oberlin, OH, USA). Muscles were accessed via dorsal and ventral incisions on the left forearm, as described previously (*23, 26, 35*). In Monkey N, specific muscles were surgically identified and implant locations were selected based on intraoperative stimulation. With Monkey R, specific muscles were surgically identified and nerve entry points to the muscles were identified (Ward et al. SFN. 2023). Nerve entry points were targeted for implants and hand function was confirmed with intraoperative stimulation. Bipolar electrodes were inserted and sutured into the muscle belly at the target location and then tunneled to an interscapular exit site. Monkey N was implanted with eight bipolar electrodes and Monkey R was implanted with 16 bipolar electrodes. Muscles targeted and evoked movements from intraoperative stimulation are described in Supplemental Table 1. Electrodes are labeled based on the muscle that they are implanted in and their expected function. Monkey N had been previously implanted (22 months prior to FES electrode implant) with two 64 channel Utah microelectrode arrays (Blackrock Neurotech, Salt Lake City, UT, USA) in the right hemisphere, targeting hand area of precentral gyrus, as described previously (*23*). Notably, these arrays were implanted 1577 days before the first BMI experiments presented here.

### Measuring joint angles

Joint angles were ultimately measured with four different methods. In both hand control and closed-loop FES experiments, joint angles were recorded as calibrated sensor values ranging from 0, fully extended, to 1, fully flexed. Sensors were calibrated to either a comfortable range of movement during hand control or to the range of movement over which FES patterns moved the hand. In hand control prior to BMI experiments, the monkeys moved their hand in a custom manipulandum (Supplemental Figure 4A) while finger (all fingers grouped together) and wrist flexion were tracked with potentiometers in the manipulandum (RH24PC, P3 America, Leander, TX, USA). The potentiometers connected to an Arduino Uno (Arduino, Somerville, MA, USA), which digitized the analog voltages and sent serial input to the xPC system. In closed loop FES experiments, due to the nerve block inactivating finger intrinsic muscles, the hand lacked rigidity and would tend to come out of the manipulandum during extension. Instead, flex sensors (FS-L-0073-103-ST, Spectra Symbol, Salt Lake City, UT, USA) were taped to the hand (Supplemental Figure 4B). These flex sensors connected to a custom analog to digital converter circuit which sent serial input to the xPC system. During open-loop experiments, such as testing stimulation patterns, movements were video recorded using a Canon Eos Rebel T3 SLR camera (Canon, Melville, NY). Wrist and index MCP angles were then extracted from video using either Deeplabcut (DLC) models (*41, 42*) and a custom MATLAB script, or directly from video frames using ImageJ (version 2.35).

### FES stimulation

Intramuscular electrodes were connected to a NNP evaluation system for stimulation. This system contained the same circuitry as the NNP used in humans (Clinical Trial NCT02329652) however was in a form more conductive to preclinical experimentation. The NNP system consisted of one power module and three pulse generator boards, each with four output channels. The return electrode of each bipolar pair connected to the same pulse generator board were electrically tied together to make one common return for each pulse generator board. All stimulation was current controlled and delivered at 32 ms inter-pulse intervals with either 5 mA or 10 mA amplitudes. Stimulation used charge-balanced biphasic pulses. Stimulation intensity was modulated by changing the width of the initial cathodic pulse, varying between 0 and 255 microseconds. Pulse width modulation was used due to the history of successful upper limb stimulation using it (*43*) and the potential to evoke the same movements with a lower amount of delivered charge (*64*).

### Designing FES stimulation patterns

Stimulation patterns were manually designed using anatomical knowledge and results from single electrode stimulation characterization. During the characterization experiments, for each monkey, the median, radial, and ulnar nerves of the monkey’s arm were blocked with a lidocaine epinephrine mix (see Nerve Block below). Then, video was obtained while each electrode was stimulated at a range of pulse widths at either 5 mA or 10 mA pulse amplitudes. Initial patterns were made by setting the pulse width at maximal commands to the pulse width that evoked the largest movement in the desired DOF without evoking substantial movements in additional DOF and the pulse width at the 50 percent command to the pulse width that caused the first movement in the desired DOF. The patterns then smoothly transitioned pulse widths between these two points. At the beginning of experiments, these established stimulation patterns were tested and then adjusted if necessary. As a result, patterns typically changed with each experiment. Example stimulation patterns for Monkey R can be found in Supplemental Figures 2A and 2B.

### Nerve Block

In order to achieve temporary hand and wrist paralysis, we performed an ultrasound guided nerve block of the median, radial, and ulnar nerves just proximal to the elbow prior to stimulation. First, lidocaine (2%) was injected subcutaneously near the planned injection sites to help prevent discomfort from injections. Then a solution of lidocaine (2%) and epinephrine (1:100 000) was injected into the perineural space surrounding the nerve under ultrasound guidance. In later experiments, the lidocaine dosing was reduced to 1% without a noticeable change in the efficacy of nerve block. Typical blocks involved injecting 1.0 mL to 1.5 mL around each of the three nerves for a total lidocaine injection of up to 8 mg/kg at 2% lidocaine, or 4 mg/kg at 1% lidocaine. Note that this maximum lidocaine dosing is smaller than similar studies which have used approximately 20 to 30 mg/kg (*36, 37*). This blocked hand function for sufficient time to perform FES tests (about two hours) without re-dosing. Block effectiveness was tested with a grasping task where we observed their motor capabilities while they attempted to grasp offered treats.

### Open*-*Loop FES Experiments

In order to investigate the range of movements that could be evoked in each DOF with FES, predetermined stimulation commands were sent, and then evoked movements were recorded on video. Experiments were performed on separate days due to muscle fatigue. These included testing stimulation at multiple points along individual patterns (four experiments with Monkey R, five with Monkey N), testing stimulation on the finger pattern with a constant wrist muscle activation maintaining a wrist posture (three experiments with each monkey), and testing stimulation at random points along both patterns (two experiments with Monkey R and four with Monkey N). Stimulation in these first two types of experiments was controlled by the experimenter. Stimulation at random points on the patterns was coordinated using xPC. The xPC selected random targets along the whole range of flexion for each DOF but did not present them on the screen. Instead, stimulation at the point in the patterns expected to reach that target were delivered and the evoked movements were recorded. Stimulation during the random sampling occurred for three seconds before changing to a new stimulation command.

### Closed-loop FES experiments

To investigate graded control of FES in 2-DOF, FES was used to move the monkey’s nerve blocked hand to acquire virtual targets. After nerve block, flex sensors were taped to the back of the wrist and index finger instead of using the manipulandum. Before closed-loop tests began, stimulation patterns for both the wrist and the fingers were tested and adjusted if necessary. Flex sensor measurements were calibrated so that the virtual hand was fully flexed or extended at approximately the range of motion available from stimulation. Target amounts of flexion for both the wrist and the fingers were selected and then targets were presented on the screen. Target sizes were 22.5% and 24.9% of the movement range for the fingers and wrist respectively. For a trial to be successful, both the fingers and the wrist had to remain in their respective targets for 500 ms.

Trials resulted in failure if this took longer than 10 seconds. The stimulation was controlled with a simple algorithm that, every 32 ms, checked if the fingers and the wrist were in their targets, and if not would update the stimulation pulse width for the joint not in the target using the stimulation patterns by moving one command value (out of 255) on the corresponding joints pattern in the direction of stimulation needed to get to the target (i.e. flexion or extension).

### BMI Experiments

To investigate whether Monkey N could use a BMI to control 2-DOF virtual wrist and finger movements before and after temporary paralysis, we implemented a position/velocity ReFIT Kalman Filter (RKF) similar to what we have previously published with two finger-groups (*23, 26*) but with updated parameters to use the wrist instead of a second finger-group. To train the decoder, first Monkey N completed at least 375 trials with hand control (average 480 per session), in the adapted finger and wrist manipulandum (Supplemental Figure 4A), while spiking band power (*65*) and finger and wrist movements were recorded. Then a position/velocity Kalman filter (KF) was trained using the hand control data and used in BMI control for at least 200 trials (average 299 trials per session). This KF BMI control data was then used as training data for the RKF model. The RKF model was then used in BMI control for at least 150 trials (average 192 trials per session) to get an initial performance, after which the nerve block was performed. Once hand function was confirmed to be lost, the same RKF model was used in BMI control again for at least 150 trials (average 186 trials per session). One more RKF model (Re-ReFIT model, ReRKF) was then trained using these post-nerve block RKF BMI control trials. The ReRKF model was then tested in BMI control for at least 125 trials (average 154 trials per session).

Targets were presented in a center-out fashion requiring flexing or extending the fingers and or wrist, for a total of eight movement combinations and to three different magnitudes of movement from center. The subsequent trial then required moving back to center, additionally the trial after a failed trial required moving back to center. Targets presented had a diameter of 16.9 to 18.8% of the movement range for the finger target and 18.7 to 20.8% of the movement range for the wrist target. The wrist target was made larger to accommodate a larger wrist visualization. Targets had to be held for 500 ms in BMI control and 750 ms in hand control. In one run of hand control trials the target hold time was mistakenly set to 500 ms instead of 750 ms. Trials were considered failed if targets were not reached and held within 10 seconds.

### Statistical analyses

Unless otherwise noted, significance for metrics such as evoked ranges of motion, and bootstrapped coefficients of determination, were tested with two-sample t-tests. Significant changes in success rate between two runs of trials was determined using a chi-squared test and significant changes in median BMI performance metrics was determined used two-sample Kolmogorov-Smirnov tests. Significant changes in BMI performance before and after nerve block across multiple sessions is tested with a two-way ANOVA where one factor is trial type (i.e. before block RKF, after block RKF, or ReRKF) and a second factor is experiment day.

## List of Supplementary Materials

Supplemental Methods

Supplemental Figures 1 to 4

Supplemental Tables 1 and 2

Supplemental Movie 1

## Acknowledgments

We thank the University of Michigan Unit for Laboratory Animal Medicine for expert veterinary and surgical support. We appreciate the support of the University of Michigan Biointerfaces Institute. We also appreciate the support from Eric Kennedy who performed the nerve block procedures and provided animal support, Jake Joseph who assisted is extracting joint angles, and Suraj Kholwadwala who assisted in initial manipulandum designs.

## Funding

National Science Foundation grant 1926576

National Science Foundation grant 2223822,

National Science Foundation Graduate Research Fellowship Program under grant 1841052

National Institutes of Health grant T32NS007222

National Institutes of Health grant R01NS105132

The Dan and Betty Kahn Foundation grant AWD011321

University of Michigan Robotics Institute

A. Alfred Taubman Medical Research Institute

## Author contributions

Conceptualization: MJM, CAC

Methodology: MJM, LHC, ETL, ALW, JWLL, MSW, NGK, TAK, PGP

Investigation: MJM, LHC, MMK, JTC, HT, DMW

Visualization: MJM, LHC

Validation: MMK

Funding acquisition: CAC, PGP

Supervision: CAC, PGP

Writing – original draft: MJM

Writing – review & editing: All authors

## Data and materials availability

Data and code will be made available upon publication.

