## Supplemental Methods, Figures, and Tables for "Functional Electrical Stimulation and Brain-Machine Interfaces for Simultaneous Control of Wrist and Finger Flexion"

### Extracting joint angles from video

During open-loop stimulation experiments, wrist and index MCP angle were extracted from video of the hand. To ensure consistent video, the camera was mounted on a tripod and angled such that the forearm just proximal to the wrist was at the edge of the frame. The monkey's forearm was braced by an experimenter to keep the hand off the table and moving within the plane of the recorded video. Deeplabcut (DLC) models (Mathis et al., 2018; Nath et al., 2019) were trained to extract hand poses from the video. To train models, 20 frames were extracted from up to three videos in an experiment. Frames were manually labeled using the DLC API and then the network model was trained. This work used the provided ResNet-50 model and initiated network training with the provided pre-trained weights. Seven points were labeled in each frame: the forearm, wrist, back of the hand, index metacarpophalangeal (MCP) joint, index proximal interphalangeal (PIP) joint, and index distal interphalangeal (DIP) joint. Due to inconsistencies in recorded video each day, this process was done for each experiment where continuous joint angles were extracted. After evaluating models on videos, frames were filtered out if model certainty was not above 95% for at least two of the labels in the frame.

Wrist and index MCP angles were extracted from DLC labels using a custom MATLAB script. An example frame with wrist and MCP angles labeled is shown in Figure 1D. The wrist angle was defined as the angle between a vector from the forearm to the wrist label and a vector from the wrist label to the MCP label. Due to commonly occurring obstructions of the forearm marker (i.e. the edge of the frame, the arm restraint, or the experimenters), the edge of the frame at the forearm was used instead of the forearm marker. The point was chosen such that the vector from the forearm to the wrist was perpendicular to the edge of the frame. The index MCP angle was defined as the angle between a vector from the wrist label to the MCP label and a vector from the MCP label to the PIP label. Extracted angles were then smoothed with a Gaussian-weighted moving average filter with a 0.5 second window. In open-loop tests where continuous joint angles were not needed, single frames were extracted from videos and joint angles were measured using ImageJ (version 2.35).

**Range of Motion Analyses**

Finger and wrist ranges of motion are calculated as the difference in index MCP joint flexion or wrist joint flexion, respectively, between a frame when stimulation is flexing the fingers or wrist and when stimulation is extending the fingers or wrist. One finger range and one wrist range was obtained for both the wrist and finger pattern tested on one experimental day. When testing the 2-DOF movement ranges, evoked movements are calibrated to the maximum angles that an individually tested pattern moved the joint to. For example, a wrist extension of 1 would be equivalent to the angle that the wrist pattern extended the wrist to on that day, whereas a finger extension of 1.0 would be equivalent to the angle that the finger pattern extended to fingers to. In one experiment with Monkey N the wrist pattern was not tested initially and the wrist movements are calibrated to the movements due to the finger pattern. Calibrated angles are then plotted into the two-DOF wrist and finger extension space. A boundary is calculated around all of the angles using the MATLAB boundary function with a shrink factor of 0.5 which balances finding a convex hull boundary and a compact boundary around the set of points.

When calculating the goodness of fit between stimulation and the evoked postures during the random stimulation sampling, one wrist and finger posture was calculated per stimulation by averaging the posture in the last 0.5 seconds of the three second stimulation. A linear regression, which included a bias term, was then fit between the stimulation command and the evoked posture.

A separate model was fit between each stimulation command, or both stimulation commands, and either the wrist or finger posture and then a coefficient of determination was calculated. A bootstrap method was used to estimate a variance on the coefficient of determination. Each model fitting was performed 100 times using n stimulation trials sampled with replacement, where n was the number of three second stimulation trials performed.

**Neural Features**

Motor-related spiking band power (SBP) (Nason et al., 2020) was recorded from 96 channels of Monkey N's Utah arrays in primary motor cortex. 96 channels were used due to available recording hardware. Briefly, we configured the Cerebus Neural Signal Processor (Blackrock Neurotech) to sample signals at 2000 Hz and band-pass filter the recorded signals between 300-1000 Hz. Continuous neural data was sent to the xPC computer which calculated a mean absolute value in non-overlapping 32-ms bins for decoding. Neural channels were masked to only use channels that were not saturated with noise and that contained morphological spikes during the experiment or in the recent past.

**BMI Decoder Training**

Decoder training matches training for a KF for two finger groups (Nason et al., 2021) but with updated parameters to use the wrist instead of a second finger group. The KF and RKF assume a kinematic state with one position and velocity for each degree of freedom. For the 2D finger and wrist task, there is one position and velocity for the wrist and one position and velocity for the fingers:

$$x_t = \begin{bmatrix} P_I \\ P_W \\ V_I \\ V_W \\ 1 \end{bmatrix}$$

Where $x_t$ is the kinematic state at time t, $P_I$ and $V_I$ are position and velocity for the index finger, and $P_W$ and $V_W$ are position and velocity for the wrist. Note that the fingers were grouped together in visualizations by giving the rest of the fingers the same kinematics as the index finger. The KF updated the kinematic state at 32 ms intervals by combining a prediction of the state given the previous state with a state update gained from the observation model which relates neural activity to the kinematic state.

The RKF and ReRKF models followed the same KF algorithm but used retrained model parameters. To train these models, the predicted velocities from the BMI trials used as training data were augmented by identifying time steps with velocities in a direction away from the target and rotating the velocities to point towards the correct 2D target vector, but not rescaling the vector. This matches the assumption that the monkey intended to be moving towards the target during the online trials. New model parameters were then trained using the augmented velocities. During use of the RKF and ReRKF models, there was no further modification of the velocities.

**BMI Performance Metrics**

We measured performance during FES closed-loop trials and BMI trials with success rate and acquisition time. In FES control, trials were excluded if both DOF started in their target, or if the taped-on flex sensors were noted as being out of place and giving erroneous readings. Success rate was then measured as the ratio of successful FES or BMI trials to the number of valid FES or BMI trials. Acquisition time was measured as the time required to complete the trial, minus the hold time and was only measured for valid successful trials. In BMI trials, we also calculated an orbiting time, that is the time between first reaching both targets simultaneously, and successfully holding the targets, minus the hold time. Orbiting time was measured for all valid and successful trials with a non-zero orbiting time.

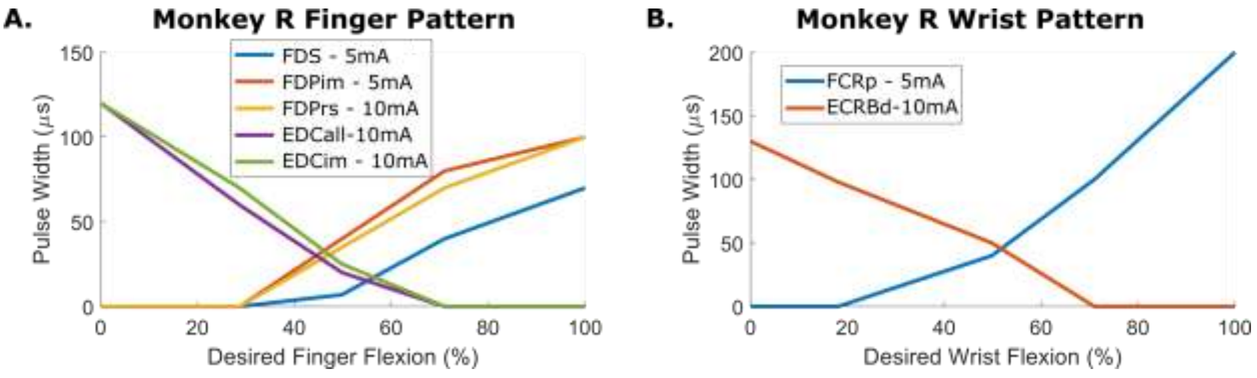


**Supplemental Figure 1 – Example stimulation patterns.**

(A) Stimulation pattern used to control the finger DOF in one session with Monkey R. The stimulation commands from 0-255 were mapped directly to the desired flexion. (B) Example stimulation pattern used to control the wrist DOF in the same session as (A) with Monkey R.

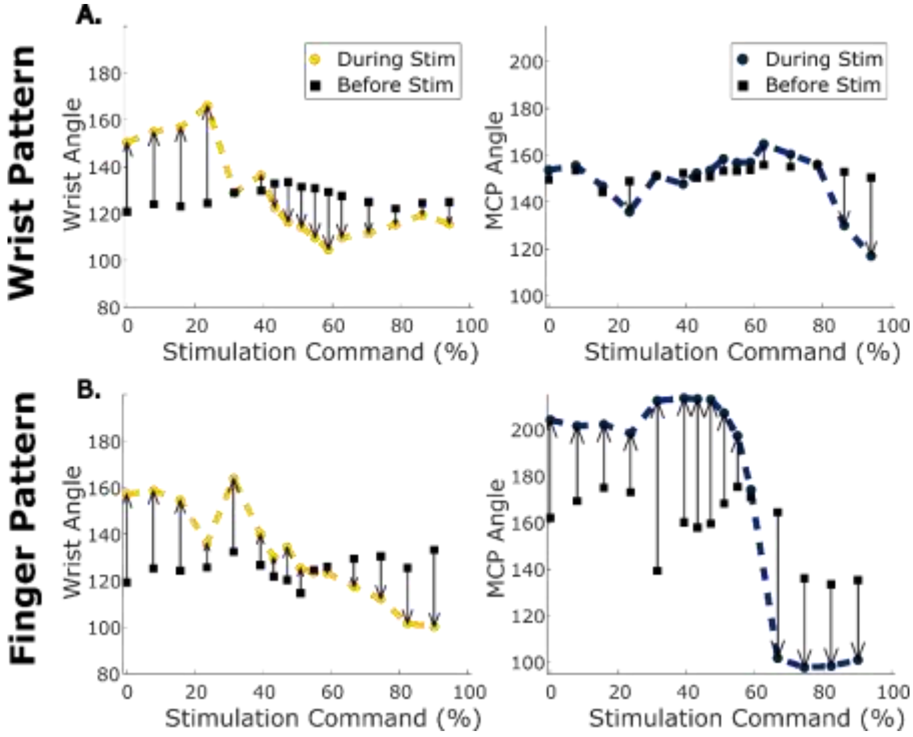


**Supplemental Figure 1 – Example measurements from testing stimulation throughout the wrist and finger pattern. (A)** Measured wrist (left, yellow) and index MCP (right, blue) angles from testing the wrist stimulation pattern on this day with Monkey R. Squares indicate the resting position before the stimulation started. **(B)** Same as (A) but for the finger stimulation pattern.

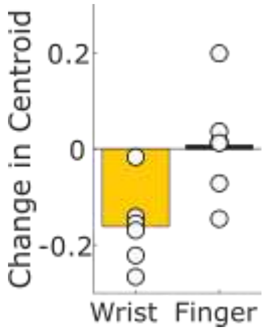


**Supplemental Figure 3 – Change in range of motion centroid between early and late trials.** Change in centroid between the boundary of evoked postures (as in Figure 4D) during the first 3 minutes and last 3 minutes using two days for monkey R and four days for monkey N.

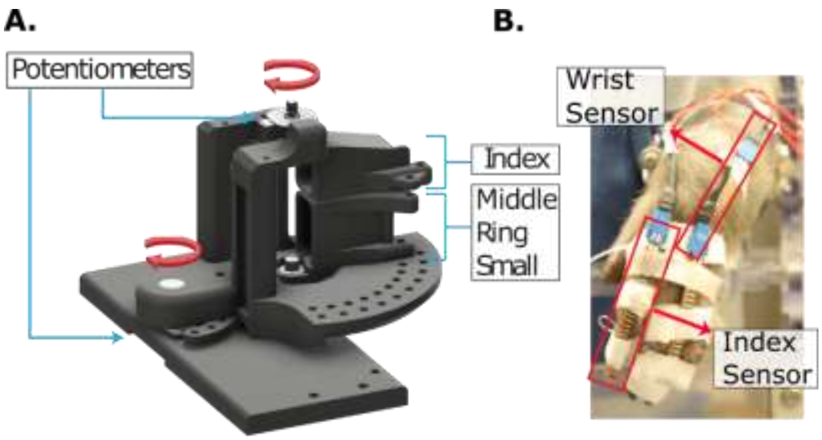


**Supplemental Figure 4 - Additional finger and wrist angle measurement methods.** (A) Illustration of the manipulandum used to measure wrist and finger flexion during BMI experiments. (B) Picture of bend sensors taped onto the monkey's hand during FES experiments. Bend sensors were used during closed loop FES experiments.

| Monkey R Electrodes | Expected Function |
|---|---|
| Flexor Digitorum Superficialis (FDS) | All finger flexion |
| Flexor Digitorum Profundus, Ulnar Site (FDPr) | Ring finger flexion |
| Flexor Digitorum Profundus Radial Site (FDPi) | Index finger flexion |
| Flexor Digitorum Profundus Ulnar Site (FDPrs) | Ring and small finger flexion |
| Flexor Carpi Radialis Proximal Site (FCRp) | Wrist flexion |
| Flexor Carpi Radialis Distal Site (FCRd) | Wrist flexion |
| Flexor Carpi Ulnaris Proximal Site (FCUp) | Wrist flexion and adduction |
| Flexor Carpi Ulnaris Distal Site (FCUd) | Wrist flexion and adduction |
| Extensor Digitorum Communis Ulnar Site (EDCrs) | Ring and small finger extension |
| Extensor Digitorum Communis Radial Site (EDCi) | Index finger extension |
| Extensor Digitorum Communis Middle Site (EDCall) | Middle, ring, and small finger extension |
| Extensor Digitorum Communis Proximal Site (EDCim) | Index and middle finger extension |
| Extensor Indicis Proprius (EIP) | Index finger extension |
| Extensor Carpi Radialis Brevis Proximal Site (ECRBp) | Wrist extension |
| Extensor Carpi Radialis Brevis Distal Site (ECRBd) | Wrist extension |
| Extensor Carpi Ulnaris (ECU) | Wrist ulnar deviation |
| **Monkey N Electrodes** | **Expected Function** |
| Extensor Indicis Proprius (EIP) | Index finger extension |
| Flexor Digitorum Profundus, targeting MRS (FDP) | Middle, ring, and small finger flexion |
| Extensor Digitorum Communis (EDC) | All finger extension |
| Extensor Carpi Radialis Brevis (ECRB) | Wrist extension |
| Flexor Carpi Ulnaris (FCU) | Wrist flexion and adduction |
| Flexor Carpi Radialis (FCR) | Wrist flexion |
| Flexor Digitorum Profundus radial proximal site (FDPip) | Index finger flexion |
| Flexor Digitorum Profundus radial distal site (FDPid) | Index finger flexion |

**Supplemental Table 1 – List of implanted electrodes and expected function.**

| | Number of successful trials | Success Rate (%) | Acquisition Time (ms) | Time to Target (ms) | Orbit Rate (%) | Orbit Time (ms) |
|---|---|---|---|---|---|---|
| Session 1 | | | | | | |
| Hand Control | 583 | 96.2 | 804 | 570 | 48.4 | 473 |
| KF Control | 353 | 97.8 | 1131 | 683 | 56.7 | 864 |
| RKF Control (Pre-block) | 257 | 97.7 | 1195 | 683 | 59.9 | 784 |
| RKF Control (Post-block) | 135 | 84.9 | 1387 | 1067 | 43.0 | 624 |
| ReRKF Control | 159 | 94.6 | 907 | 779 | 32.1 | 736 |
| Session 2 | | | | | | |
| Hand Control | 375 | 94.0 | 1073 | 695 | 41.3 | 1350 |
| KF Control | 282 | 95.3 | 1131 | 811 | 47.5 | 768 |
| RKF Control (Pre-block) | 145 | 98.0 | 971 | 843 | 37.2 | 912 |
| RKF Control (Post-block) | 186 | 93.0 | 1611 | 939 | 55.4 | 1152 |
| ReRKF Control | 125 | 96.9 | 1003 | 715 | 52.0 | 512 |
| Session 3 | | | | | | |
| Hand Control | 431 | 99.1 | 738 | 581 | 27.8 | 679 |
| KF Control | 228 | 94.6 | 1522 | 859 | 60.1 | 1184 |
| RKF Control (Pre-block) | 143 | 86.1 | 1227 | 779 | 51.7 | 768 |
| RKF Control (Post-block) | 166 | 83.0 | 1650 | 1074 | 51.8 | 1040 |
| ReRKF Control | 156 | 94.0 | 1563 | 843 | 58.3 | 1376 |

**Supplemental Table 2 – Performance metrics during all BMI control experiments**

**Supplemental Movie 1 – Open loop control of finger movements in different wrist postures.**
